# Feeding a low-protein diet exacerbates the susceptibility to Citrobacter rodentium and Dextran Sulfate Sodium-induced intestinal injury in mice

**DOI:** 10.1101/2025.10.12.681925

**Authors:** Mehakpreet K. Thind, Mary Flanagan, Sanne de Wit, Aida Glembocki, Jiali Pan, Celine Cuinat, Sharon Leung, Peter Kim, Elena M. Comelli, Amber Farooqui, Celine Bourdon, Robert H. J. Bandsma

## Abstract

Globally, almost half of all early childhood deaths are linked to severe undernutrition, herein referred to as severe malnutrition. Mortality in severely malnourished children is often attributed to common infectious diseases, including enteric infections. It has been proposed that impaired intestinal barrier function contributes to mortality, but direct evidence is limited and thus there exists a need to develop improved preclinical models to test these and other mechanistic hypotheses. In this study, we aimed to describe differences in response to enteric inflammation and infection in the colon of malnourished mice compared to well-nourished littermates. C57Bl/6 male weanlings were fed isocaloric diets, either a low 1% protein diet (LPD) or a control 18% protein diet (CPD) for 2-weeks either in combination with oral administration of dextran sodium sulfate (DSS), or *Citrobacter rodentium* (*C. rodentium)*. LPD-fed mice were more susceptible to DSS or *C. rodentium* as evidenced by increased clinical severity scores, and reaching their humane endpoints. LPD-fed mice also showed more signs of colonic dysfunction with reduced levels of tight junction proteins, higher colonic pathogen load, and increased systemic inflammation and bacterial spread. Taken together, these observations show that malnourished animals have increased susceptibility to intestinal dysfunction caused by either chemical exposure or infection. These novel preclinical models can be used to further elucidate the processes involved in enteric dysfunction in malnutrition and to test therapies to improve intestinal repair and outcomes.

## Introduction

Severe malnutrition (SM), herein referring to early childhood undernutrition, is associated with high morbidity and mortality of children under 5 years of age living in low-resource settings(1). SM encompasses two major forms of malnutrition, severe wasting (most common, also known as marasmus) or nutritional edema (also known as kwashiorkor), and SM is associated with immunodeficiency and environmental enteropathy (EE) in children(2–4). EE is characterized by high exposure to environmental microbes and linked to villous atrophy, intestinal inflammation and signs of epithelial barrier dysfunction (5–8), which is strongly related to poor clinical outcomes(7–11). In fact, in children with SM and EE, exposure to common pathogens can become life-threatening(2,4,8,10–15). Gram-negative bacteria commonly found in blood of acutely ill children with SM and associated with diarrhea include enteric pathogens such as *Salmonella typhimurium*, *Shigella spp,* enteropathogenic *Escherichia coli* (EPEC), and *Klebsiella pneumoniae*(16–19). This suggests that children with SM are more vulnerable to bacterial translocation across the intestinal lumen leading to sepsis. This could be more likely to occur in the large intestine where bacterial density is the highest. In addition, there exists a positive association between diarrhea and mortality in children with SM(17,20). Despite following established treatment regimens, including the widespread use of broad-spectrum antibiotics, the in-patient and post-discharge mortality rate of children with SM remains high(1) and therefore should be a prioritized area of research.

Pre-clinical models can be used to elucidate the pathophysiological processes and identify treatment approaches that could improve intestinal dysfunction associated with SM (21–24). This would otherwise not be possible as non-invasive methods in this clinical setting are limited and invasive methods such as endoscopy and colonoscopy pose a risk to this highly vulnerable population. Several animal models have been developed to study malnutrition, typically wasting, and intestinal changes(5,6,15,22,23,25–28). For example, mice fed a moderately low-protein diet (7%) developed enteric dysfunction if also exposed to specific bacteria (i.e., Bacteroidales and *E. coli* strains) (15). Similarly, post-weaning mice fed a moderately-low protein diet (5.8%) developed intestinal dysfunction after receiving an additional gastrointestinal insult (indomethacin or lipopolysaccharide (LPS))(29). In these models, wasting was not severe and, while intestinal dysfunction was induced after the additional insult, the mice did not develop severe infection.

More recently, a study showed that feeding a corn-based diet with nutritional deficiencies in protein, fat, minerals and vitamins, as commonly consumed in resource poor settings, to weanling mice for 28-days was sufficient to induce anthropometric deficits, dysbiosis, enteric dysfunction and immune alterations(22). Similarly, 14-days of severe protein deficiency (1%) alone has been shown to induce highly significant linear growth impairment and wasting as well as enteric dysfunction in mice(5,6,25).

However, the susceptibility to enteric infection and inflammation in severely wasted mice, particularly in the colon, remains poorly studied. Furthermore, whether malnutrition exacerbates bacterial translocation across the intestinal barrier in the context of enteric infection has not been demonstrated directly. Dextran sodium sulfate (DSS), a chemical insult, is well-known to induce acute colitis characterized by bloody diarrhea, mucosal ulcerations and innate immune cell infiltration through its toxic effects to gut epithelial cells that impacts the integrity of the mucosal barrier, and causes subsequent colonic inflammation(30,30). *Citrobacter rodentium* (*C. rodentium)* is an extracellular, gram-negative enteric pathogen that is commonly used to model human intestinal diseases, such as EPEC and enterohemorrhagic *Escherichia coli* (EHEC) infections, in mice(31,32). It establishes colonization by adhering and forming attaching-and-effacing lesions on the colonic gut epithelium inducing an infection response similar to EPEC and EHEC(31), including crypt hyperplasia. Colonic crypt hyperplasia involves the proliferation of transit amplifying cells and is a typical tissue response to *C. rodentium* colonization(31,33–35). This current study aimed to describe novel mouse models of severe wasting and large intestinal infection and inflammation and assess the impact on intestinal epithelial barrier function.

## Results

### Low-protein diet feeding exacerbates the effect of DSS-treatment in weaning mice

First, we used DSS to induce acute colitis in mice(30) fed either a low- or control- protein diet to study differences in disease burden. Weanling C57Bl/6 male mice were randomised into two groups fed either the low 1% protein diet (LPD) or the 18% control protein diet (CPD) for 14 days. On day 7, groups were provided water containing either 1% or 2% DSS for either 3 or 5 days, respectively (Figure 1a; Figure S1a). As previously demonstrated(5,6,25), CPD-fed mice gained approximately 40% of their body weight over the 14 days, whereas LPD-fed mice lost approximately 20% of their body weight (Figure 1b; Figure S1b). The malnourished mice showed minimal clinical features and had higher clinical severity scores largely due to weight loss. When exposed to 1% or 2% DSS, the LPD-fed mice lost significantly more weight compared to controls (Figure 1b; Figure S1b). Furthermore, LPD-fed mice exposed to DSS also exhibited more clinical symptoms especially relating to diarrhea, bloody stools and reduced mobility. Thus, the DSS-exposed LPD-fed mice had higher clinical severity scores (Supplementary Table 1; Figure 1c; Figure S1c). 60% of these DSS-exposed malnourished mice died or reached pre-defined humane endpoints (Figure 1d; Figure S1d), while all DSS-exposed controls survived. Overall, these findings highlight that LPD feeding increases the host susceptibility to DSS-induced colitis.

**Figure 1:**
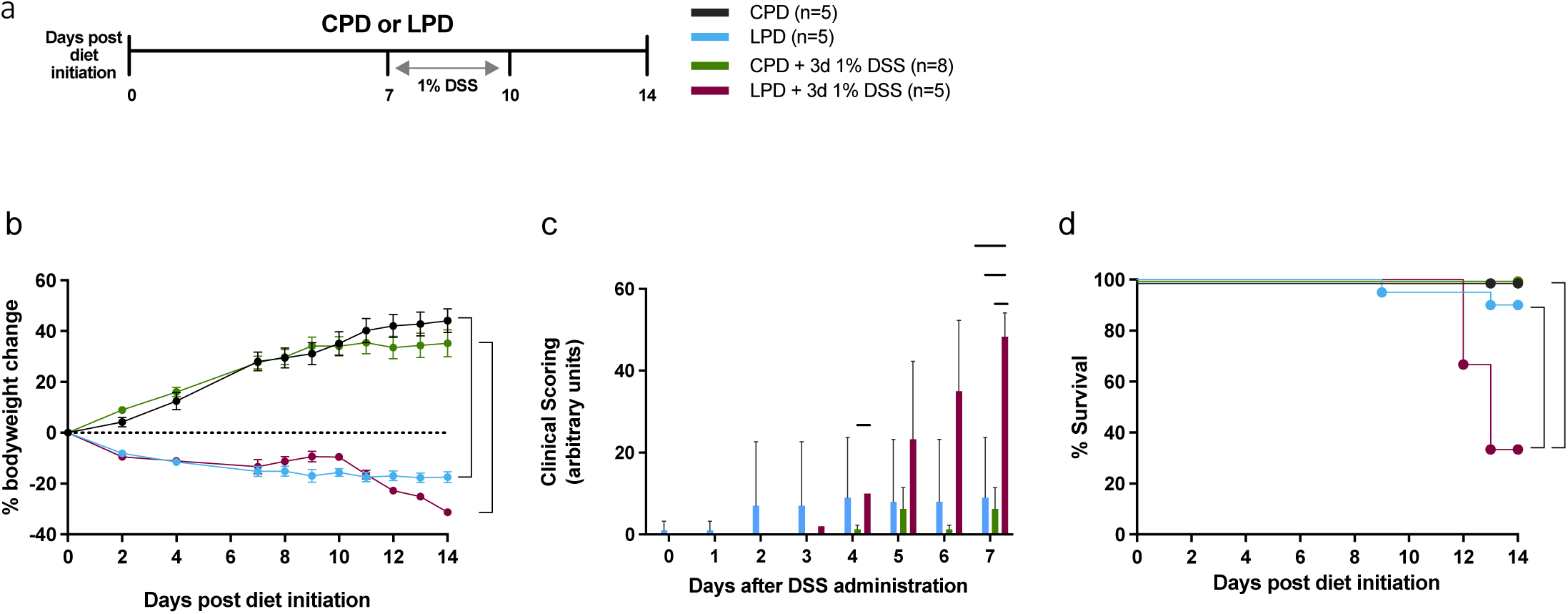
Mice fed a low-protein diet are more susceptible to DSS-induced colonic inflammation. (a) Experimental design of the DSS-induced colitis; weanling C57Bl/6 mice fed control (CPD) or the low-protein diet (LPD) were given either 1% DSS dissolved in drinking water for 3 days or DSS-free drinking water. (b) Percent bodyweight change from day 0 and (c) Clinical Severity Scoring (CSS) over time. CSS is a measurement of hunched posture, ruffled fur, immobility, weight loss, diarrhea and bloody stools(33,82). (d) Survival of CPD and LPD- fed mice. Results are expressed as means ± SD and *p < 0.05; ****p < 0.0001, as determined by (b) two-way ANOVA, (c) linear mixed-effect analysis and (d) two-sided log-rank (Mantel-Cox).

### Susceptibility to infection with *C.rodentium* is increased in low-protein diet fed mice

Next, we explored whether mice fed a low-protein diet were more susceptible to intestinal infection by *C.rodentium*, a gram-negative pathogen. To do this, we inoculated the mice by oral gavage with 10^7^ colony forming units (CFU), a lower infective dose compared to the more typically used 10^8^-10^9^ CFU (Figure 2a). This lower dose reduces the likelihood of early mortality or reaching humane endpoints prematurely in young mice(36,37). As expected(5,6,15,25,38), LPD-fed mice inoculated with *C. rodentium* had shorter tail length (which represents stunted growth) (Figure S2a). Their colon length was also shorter (Figure S2b; 5.58 ± 0.58 vs. 4.47 ± 0.45, p<0.0001) and their ratio of colon over tail length was also significantly lower compared to CPD-fed mice post-infection (Figure S2c; 0.77 ± 0.09 vs. 0.71 ± 0.06, p<0.0001). After infection, the LPD-fed mice had, at day 4 and 7, more *C. rodentium* in their stool compared with CPD mice (Figure 2b). This suggests that malnourished mice experience higher pathogen colonization and/or have a reduced ability to clear pathogenic bacteria. Similarly, systemic levels of IL-22, a pro-inflammatory cytokine, were higher in LPD-fed mice 7 days post- *C. rodentium* infection (Figure 2c). With infection, 25% of LPD-fed mice reached the humane endpoint, mostly due to more than 25% weight loss (Figure 2d-e). Reaching their humane endpoint prevented us from studying mice for a longer period post-infection. As previously reported(39,40), no change in body weight was observed in CPD-fed mice after *C. rodentium* infection.

**Figure 2:**
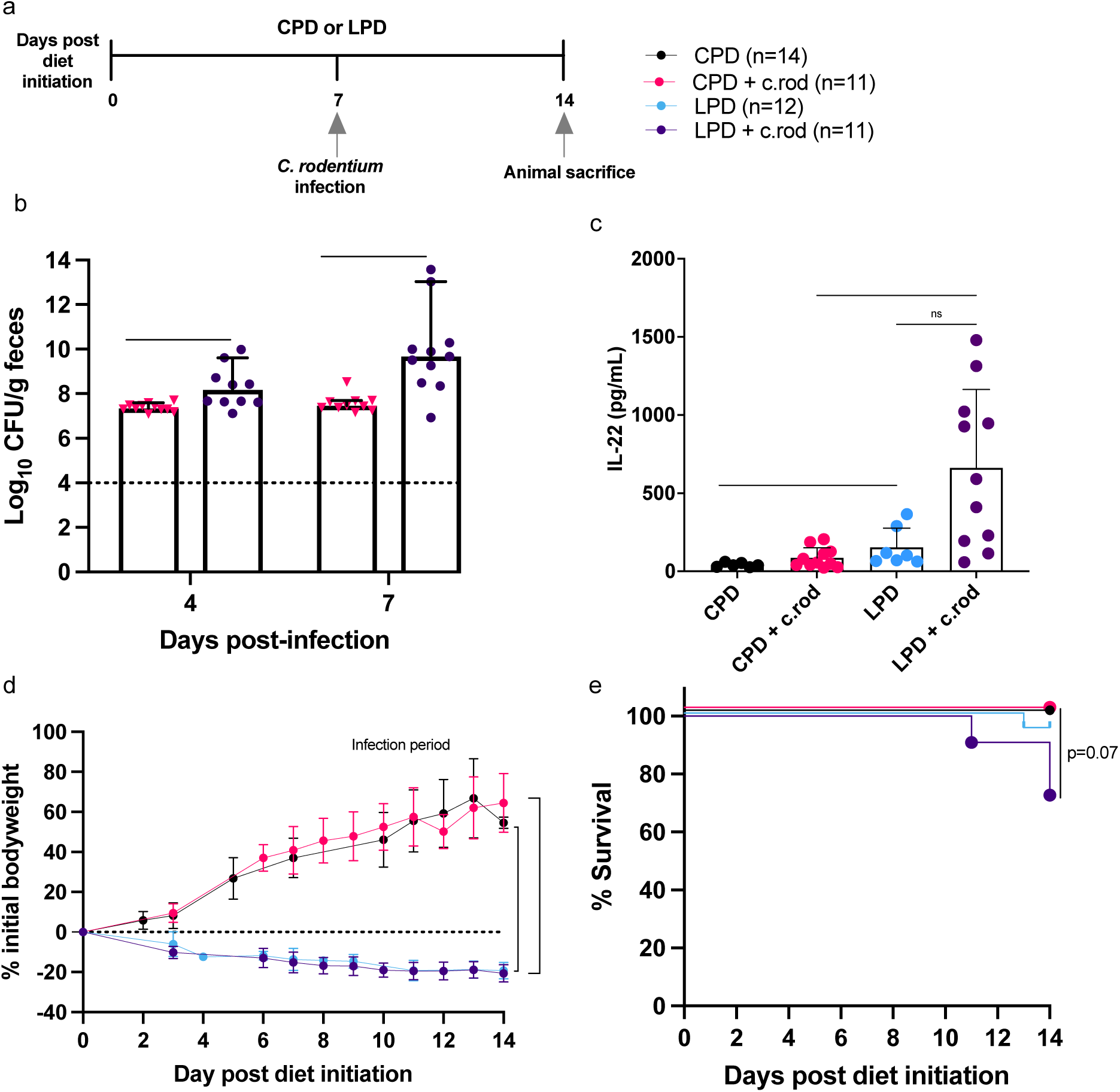
Mice fed a low-protein diet are more susceptible to *Citrobacter rodentium* infection. (a) Experimental design of *Citrobacter rodentium* (*C. rodentium*) infection; weanling C57Bl/6 mice fed control (CPD) or the low-protein diet (LPD) were inoculated by gavage with 1X10^7^ colony forming unit (CFU) of *C. rodentium*. (b) qPCR quantification of *C. rodentium* in stool at day 4 and 7 post-infection in CPD and LPD mice. Dotted line represents the limit of detection for *C. rodentium*, as previously published(83,86). (c) Mice were sacrificed on day 7 post-infection and IL-22 concentration was measured by ELISA in serum. (d) Change in body weight and (e) survival of CPD and LPD mice. Results are expressed as means ± SD and **p < 0.01; ***p < 0.001; ****p < 0.0001, as determined by (b) two-way ANOVA, unpaired two-tailed Mann-Whitney test, and Fisher’s exact test, (c) one-way ANOVA with Tukey’s multiple comparisons test and unpaired two tailed t-test analysis, (d) two-way ANOVA and mixed-effect analysis, and (e) two-sided log-rank (Mantel-Cox).

### Crypt hyperplasia caused by *C. rodentium* is blunted in low-protein diet fed mice

We next determined whether the higher bacterial load and lower survival of LPD mice was associated with changes in hyperplasia of colonic crypts that is typically observed in response to *C. rodentium*. Without infection, LPD-fed mice had a crypt length that did not differ from controls (Figure 3a-b). Also, the colons of LPD-fed mice did not show differences in histological signs of inflammation (Figure 3c). CPD-fed mice infected with *C. rodentium* trended towards increased colonic inflammation and significant crypt hyperplasia as observed by increased crypt length (Figure 3a-c). This typical response to *C. rodentium* is thought to prevent systemic pathogen spread, and promote repair of the epithelium(31). In contrast, infected LPD-fed mice showed little crypt hyperplasia (Figure 3a-b), and insignificant colonic inflammation compared to LPD-alone and infected CPD-fed mice (Figure 3c). In line with this, the expression of Ki67, a marker of cell proliferation, trended towards an increase in the colonic crypts of CPD-fed mice but its expression was unaltered in the colonic crypts of LPD-fed mice post-infection (Figure 4). These findings suggest that malnourished mice have a reduced ability to initiate a protective colonic response to *C. rodentium,* potentially leading to poor pathogen clearance.

**Figure 3:**
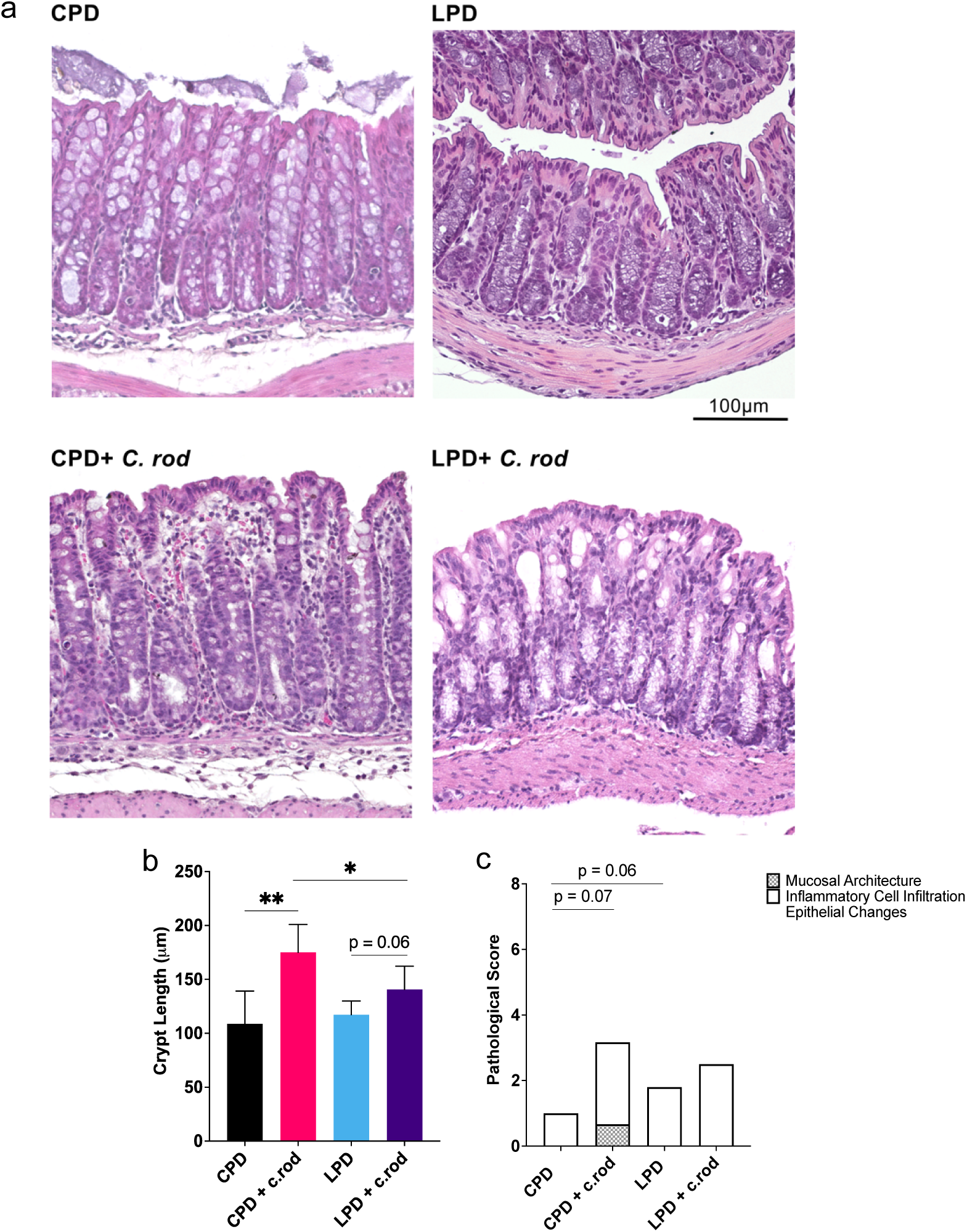
Assessment of colonic inflammation due to *Citrobacter rodentium* infection. (a) Representative images of colon (H&E, 20x). Scale bar, 100 µm. (b) Quantification of crypt hyperplasia calculated from the mean length of 30 crypts per animal (n ≥ 5/group) and (c) and cumulative pathology score on day 7 post infection (n ≥ 5/group). Results are expressed as means ± SD; *p < 0.05; **p < 0.01, as determined by (b-c) one-way ANOVA with Tukey’s multiple comparisons test and unpaired two tailed t-test analysis.

**Figure 4:**
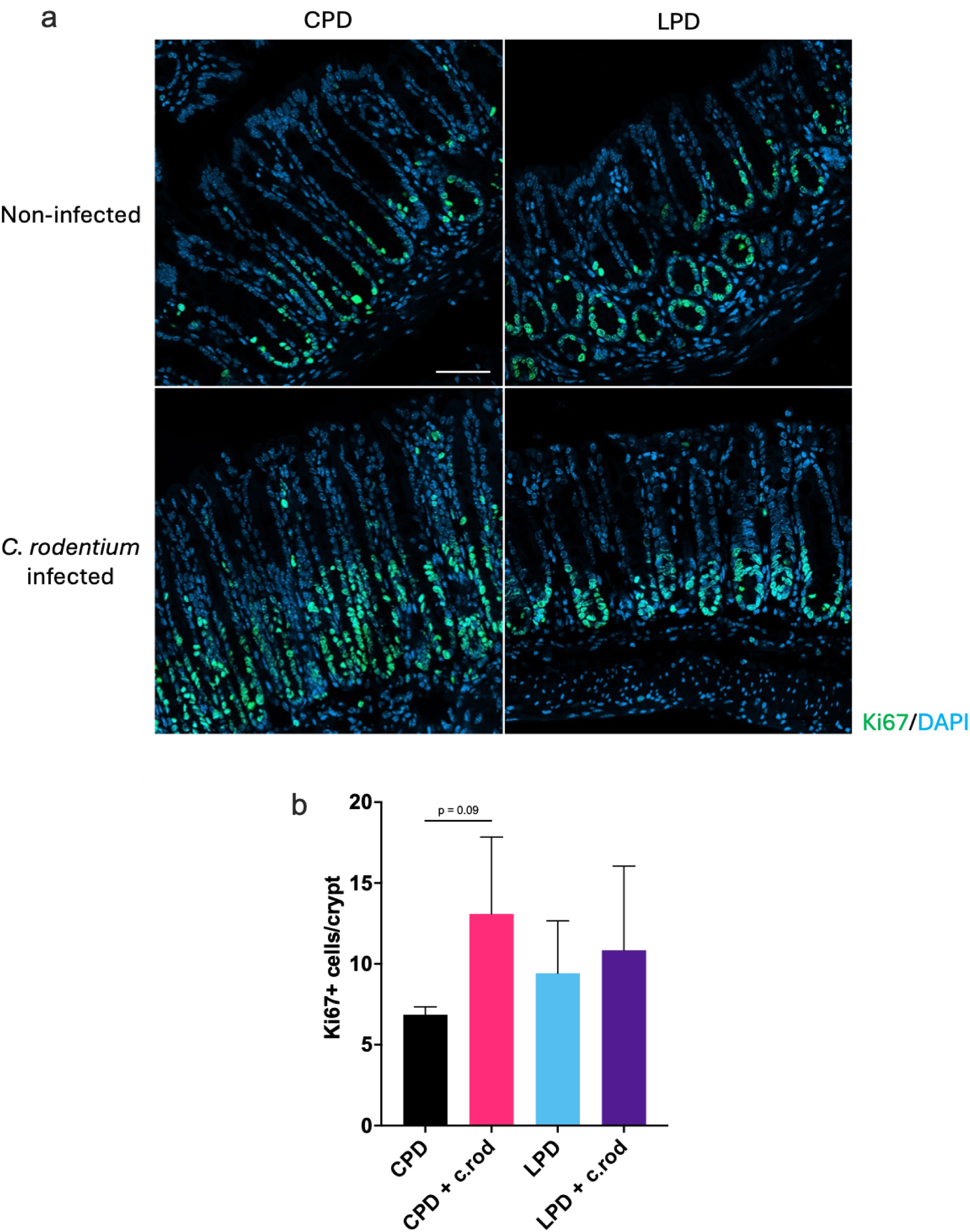
Expression of Ki67, a proliferation marker, in colon of *Citrobacter rodentium* infected mice. (a) Representative confocal microscopy images (63x) of proliferation marker Ki67 (green) in the colon with DAPI counterstaining of nuclei in blue. Scale bar, 50 µm. (b) Quantification of Ki67+ cells per crypt in the colon (n ≥ 3 mice, 4 crypts per mouse). Results are expressed as means ± SD, as determined by (b) one-way ANOVA with Tukey’s multiple comparisons test and unpaired two tailed t-test analysis.

### Increased susceptibility to *C. rodentium* is associated with loss of colonic barrier integrity in low protein diet-fed mice

To assess whether *C. rodentium* infection and invasiveness was associated with loss in colonic barrier integrity, we measured the tight junction proteins CLD-3 and occludin. In uninfected LPD-fed mice, changes in CLD-3 were not observed (Figure 5a-b), but occludin levels were higher compared to controls (Figure 5a,c). With *C. rodentium* infection, tight junction proteins differed between diet groups, where the colonic expression of CLD-3 decreased in LPD-fed mice compared to controls (Figure 5a-b). However, occludin was similarly elevated in all conditions compared to CPD mice (Figure 5a, c). Given the impact of *C. rodentium* on barrier integrity in this study, we evaluated for evidence of bacterial translocation in the spleen and liver, tissues known to be first exposed to bacteria entering the circulation due to breaches in intestinal barrier(41–43). We found that homogenates of spleen and liver from LPD-fed mice had higher bacterial counts of *C. rodentium*, consistent with increased translocation (Figure 5d-e). Together, these findings demonstrate that LPD-fed mice show an increased susceptibility to a low infectious dose of *C. rodentium* which is associated with reduced markers of colonic barrier integrity and with higher dissemination of bacteria within the host.

**Figure 5:**
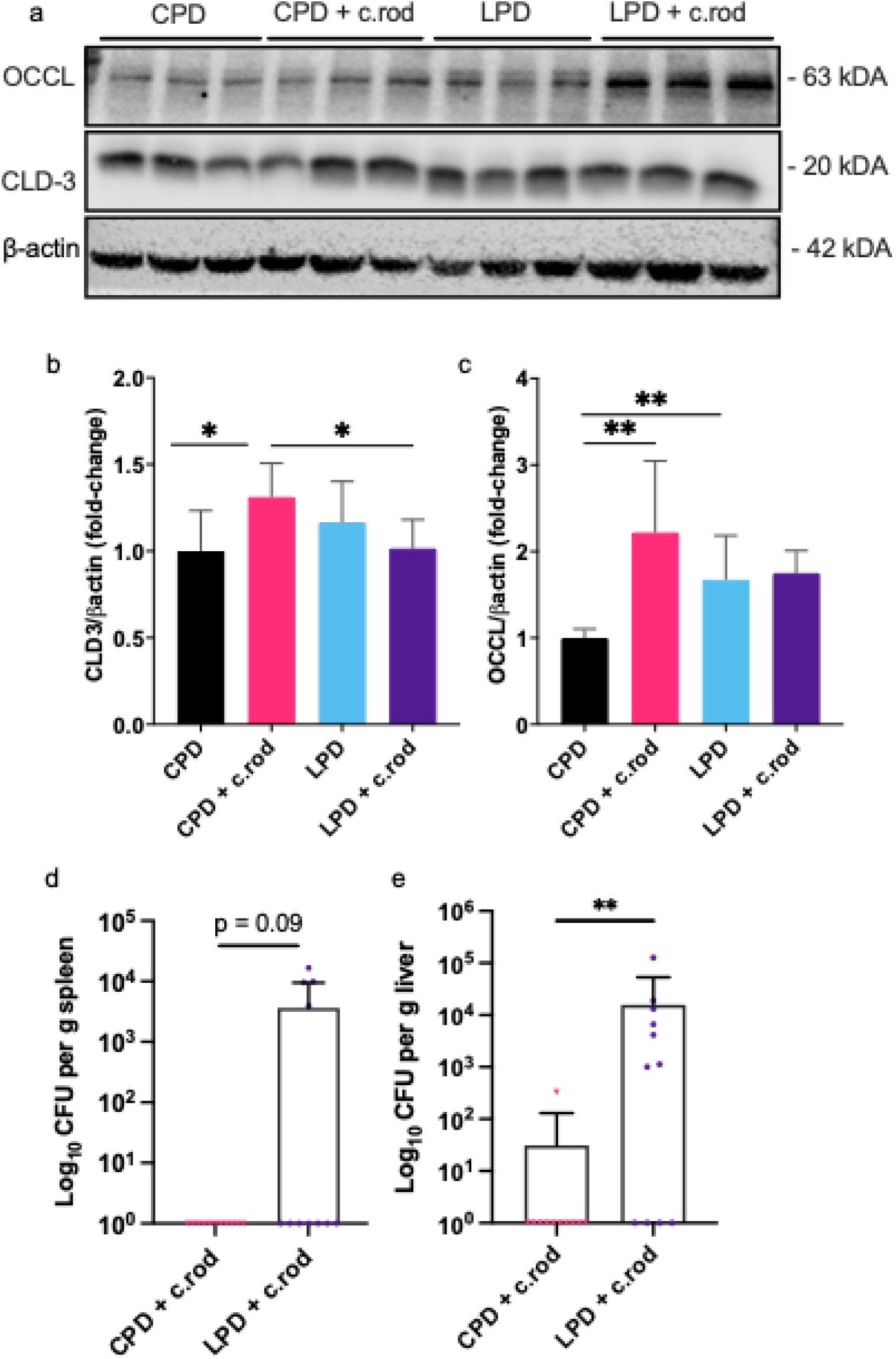
*Citrobacter rodentium* infection alters colonic structural integrity and promotes systemic bacterial translocation in malnourished mice. (a) Representative cropped western blot membranes probed for claudin-3 (CLD-3), and Occludin (OCCL) and β-actin in colonic tissue of uninfected and infected CPD and LPD-fed mice. Full uncropped blots are presented in supplementary Figure S3. Quantification of (b) CLD-3, and (c) OCCL normalized to β-actin in CPD colonic tissue was calculated (n=6/group). Bacterial burden in the (d) spleen, and (e) liver expressed as CFU per gram of tissue. Results are presented as means ± SD; *p < 0.05; **p < 0.01, as determined by (b-c) one-way ANOVA with Tukey’s multiple comparisons test and unpaired two tailed t-test analysis, and (d-e) unpaired two tailed Mann-Whitney test and Fisher’s exact test.

## Discussion

We developed two complementary novel “double insult” models of malnutrition with colonic infection/inflammation and demonstrated that mice fed a low-protein diet exhibited decreased survival and increased colonic dysfunction when exposed to chemically induced intestinal inflammation by DSS or a low dose infection by *C. rodentium*. These pre-clinical models provide valuable tools to study the pathophysiological processes underlying increased susceptibility to intestinal disease associated with malnutrition. By establishing these models, we aimed to facilitate the testing and development of novel therapeutic approaches that could improve disease response and/or accelerate tissue repair of the colon in malnourished hosts.

To our knowledge, we are the first to show that mice fed a low-protein diet exhibit an altered colonic response to exposure of DSS or *C. rodentium* compared to mice fed a regular diet. Our findings are complimentary to previous observations in the small intestine, where dietary insults alone(5,6,22) or in combination with specific bacteria(15,21,29,44) have been shown to recapitulate elements of enteropathy associated with malnutrition(22,45–48). Mice fed with a moderately low protein diet (7%) showed villous blunting, reduced barrier function (reduction of CLD4 expression) and intestinal permeability, assessed by higher fluorescein isothiocyanate (FITC)-labelled dextran in the serum. Here, dissemination of bacteria and systemic inflammation occurred only after exposure to a mixture of Bacteroidales and *E. coli*(15). Similarly, mice fed a moderately low protein (5.8%) with repeated oral doses of indomethacin also showed impaired gut barrier function (higher FITC in serum)(29).

We chose DSS and *C. rodentium* to mimic symptoms of diarrhea described in children(16,49,50). More specifically, *Shigella*, heat-stable enterotoxin-producing *Escherichia coli* (ST-ETEC), *Campylobacter*, and typical enteropathogenic *E coli* were recently shown to be the leading causes of diarrhea(51,52) and these bacterial pathogens were associated with linear growth faltering(51) and death(52). C*. rodentium* is a murine mucosal pathogen that shares pathogenic mechanisms with human EPEC and EHEC infections and has been widely used to study the pathogenesis of these two clinically important human gastrointestinal pathogens(31,53). The DSS-induced inflammatory insult is also a commonly used pre-clinical model of colitis and IBD, and this model is relevant to malnutrition since it presents with several overlapping features, such as intestinal inflammation, poor intestinal absorption, and disrupted barrier function(54,55). Despite using lower doses of DSS and *C. rodentium*, we found the models to be robust and reproducible.

Colonic hyperplasia in *C. rodentium* infection has been attributed to hyperproliferation of the transit amplifying cells and LGR5^+^ cells via activation of the Wnt/β-catenin signaling pathway(56). This increases cell turnover, facilitating shedding of infected cells into the lumen, promoting pathogen clearance, and restoring intestinal homeostasis in a self-limiting manner (31). In this study, malnourished mice exhibited a blunted crypt hyperplasia response at 1-week post-infection with C. *rodentium*, likely due to a reduced proliferative capacity as demonstrated by the reduced Ki67-positive colonic epithelial cells in mice fed a low-protein diet (Figure 4). Interestingly, unlike the jejunum and ileum of non-infected malnourished mice where a significant decrease in Ki67 expression was reported(5), there was no difference between non-infected CPD and LPD mice in the colon in our study. Overall, impaired cell turnover and shedding could potentially be related to the increased intestinal colonization and systemic dissemination in malnourished hosts observed in this study.

It is hypothesized that malnutrition is associated with an increased susceptibility to enteric infection and inflammation due to underlying enteric dysfunction and immune defects that increases susceptibility to systemic bacterial burden and inflammation, and mortality. We report for the first time that mice fed a LPD exhibit enhanced systemic translocation of pathogens that typically cause colonic infection. Scarce data existed prior to this study in support of increased risks of bacterial translocation in malnourished conditions. Mice fed a 3% protein diet deficient in calories, zinc and iron for 28 days showed exaggerated translocation of intestinal microbiota and their products, i.e., LPS, after exposure to intradermal systemic infection, compared to control mice(57). Another dietary model to induce malnutrition (7% protein and 5% fat) led to higher bacterial burdens of S. Typhimurium in the small intestine, as well as increased colonization in the liver and spleen(15). Additionally, children in Zambia, with SM and persistent diarrhea were found to have higher damage in the small intestine—reduced epithelial surface, histological lesions and disruptions of claudin-4 and E-cadherin—and subsequent higher levels of circulating LPS, which was used as a marker of bacterial translocation(7). A higher blood concentration of 16S rRNA gene copy numbers was also observed among children with complicated SM in Uganda due to impaired gut function(58).

Epithelial barrier integrity is a dynamic process that relies on several tight junction proteins to regulate paracellular permeability and protect against pathogens(59–61). When exposed to *C. rodentium,* well-nourished mice tended to increase CLD3 and occludin, whereas this cohesive increase in tight junction proteins was blunted in malnourished mice. Brown et al. also reported reductions in these tight junction proteins in mice fed a moderately low-protein diet and exposed to a mixture of Bacteroidales and E. Coli(15). Altered expression of claudins and occludins were also implicated in poor response to EPEC infections in preclinical models of inflammatory bowel disease(15,34,57,62,63). Therefore, structural perturbations in tight junction proteins during *C. rodentium* infection(40) could also contribute to the systemic spread of bacteria to the spleen and liver in severely malnourished hosts.

The increased susceptibility of malnourished mice may also be attributed to poor immune responses(64), as has been demonstrated in recent clinical studies(14,35). Early childhood malnutrition impairs both innate and adaptive immunity reducing the capacity to clear pathogens, despite high systemic and intestinal inflammation (2,4,16,65). In particular, neutrophils—key first responders to pathogens—show impaired antimicrobial responses in malnutrition despite having elevated numbers(4,13,64,66). Neutrophils are also major producers of IL-22, a cytokine that regulates crypt hyperplasia(67) and secretion of antimicrobial proteins in response to early C. *rodentium* infection to maintain the epithelial barrier function(68), (69). In DSS-induced intestinal damage, IL-22 modulates repair of colonic epithelium(70), where exogenous supplementation of IL-22 improves response. However, persistent elevations in IL-22 levels, as we observed in this study, are also implicated in chronic inflammatory conditions through sustained immune responses and inflammation(71), and could be a driver of the exacerbated pathology observed in this study in the LPD-fed mice.

This study had several limitations. The incidence of bacterial translocation may have been underestimated since translocation was only studied at a single time point post-infection. The experimental timeline was another major limitation. A 7-day timeline post-infection is insufficient to observe the full evolution of epithelial hyperplasia, which typically peaks around 10-12 days post-infection in adult mice. Multiple pathogens are commonly detected in ill children with severe malnutrition(9), and therefore, the role of gut microbiota and mixed infections should be studied further. The regulation of tight junction proteins is complex, including post-translational modifications. As such, these processes need to be studied in-depth to gain a comprehensive understanding of the role of malnutrition and tight junction proteins for intestinal barrier permeability(6,72). Additionally, mucus production by goblet cells helps to prevent systemic dissemination of bacteria(73,74), but was not explored in this study. Indeed, a decrease in goblet cell number in both malnourished rats(75–77), pigs(78), and humans(79), as well as reduction in mucin content in animal models(75) and humans(80,81) has been described. We also did not investigate infiltration of immune cells nor those more specifically linked to IL22 secretion. Other elements of the inflammatory, metabolic and organ response, including serum albumin levels, blood glucose, and liver and kidney functions were also not examined, but have been shown to be altered in clinical and preclinical models of SM

In summary, we demonstrated that LPD-fed mice exposed to DSS, or *C. rodentium* have a heightened disease susceptibility driven by an atypical colonic response to these insults compared to CPD-fed mice. These murine models provide valuable tools to investigate the pathophysiology of enteric infections in the context of malnutrition, especially in the colon where very little has been studied till date. Future research should leverage these models to identify and evaluate novel therapeutic agents that could be tested prior to being translated into clinical trials designed to improve long-term outcomes in children with severe malnutrition.

## Methods

### Animals

All mouse experiments were approved and conducted in strict accordance with the Animal Care and Use Committee guidelines (protocol number: 1000058060) at Lab Animal Services (LAS) Facility of SickKids, Toronto. Reporting follows ARRIVE guidelines. C57Bl/6J mice sourced from The Jackson Laboratory (Bar Harbor, Maine USA) were bred and maintained as a specific pathogen free colony. Male weanlings (21 days old) were weight-matched and randomized to one of two diet groups. For 14 days, mice received either: (1) a control 18% protein diet (CPD; TD: 180483), or (2) a 1% low-protein diet (LPD; TD: 180481) made by Envigo Teklad Diets (Madison, WI). The diets were isocaloric where the decrease in casein was compensated with an increase in corn starch (diet composition detailed in Supplementary Table 1). All animals were group housed at 22℃, following a 12-h light-dark cycle and had *ad libitum* access to food and water. Reseaches were unblinded to the treatment groups.

### DSS-induced colitis

To induce colitis, mice fed either CPD or LPD were administrated dextran sulfate sodium (DSS, molecular weight 40 kDa; ThermoFisher, Massachusetts, USA) in drinking water as follows: 1% DSS for 3 consecutive days or 2% DSS for 5 consecutive days starting on day 7. After dosing, animals were returned to normal drinking water. Body weight, and food/water consumption was recorded daily. To minimize animal suffering, clinical severity scores (33,82) were determined daily by trained staff for each mice post DSS administration that included clinical signs such as hunched posture, inactivity, ruffled fur, diarrhoea and bloody stools (Supplementary Table 2). A score of 35 or higher was considered the humane endpoint, and animals were removed from the study early and euthanized immediately if they reached humane endpoints prior to completing the study. All other animals were euthanized on the last day of the experiment (study day 14). The results from the mice excluded early were used for the survival curve graphs only.

### Bacterial preparation, dosing and gavage

Wildtype *Citrobacter rodentium* strain DBS100 bacteria was grown to log phase in Luria-Bertani (LB) broth with streptomycin at 37°C for 2hr to an optical density (OD) at 600 nm of 1 (3.7X10^8^ colony-forming unit (CFU)/mL). CPD and LPD-fed mice were infected on day 7 of the experimental period through oral gavage with either 100 μL of inoculum containing a total of 10^7^ CFU bacteria in PBS or of sterile PBS and were sacrificed 7 days later. To confirm dosage, left over inoculum were serially diluted in PBS, plated on LB agar and grown aerobically at 37 °C for 24 h to verify administered CFU. Monitoring of animals was performed similarly as for mice exposed to DSS-colitis. Two mice reached the humane endpoint early and were not included in further analyses.

### Euthanasia and tissue collection

After the 14 day experimental period, mice were put under anesthesia with isoflurane and euthanized by cervical dislocation. Liver, spleen and colon were collected for the *Citrobacter rodentium* infected mice. Liver and spleen were weighed, and tail and colon length were measured.

### Quantification of *C. rodentium* in stool

DNA was extracted from feces collected at 4- and 7- days post-infection using the E.Z.N.A^TM^ Stool DNA Isolation Kit (Omega Bio-Tek, Doraville, GA, USA), as per manufacturer’s instructions. DNA concentration was quantified using a Nanodrop (Thermo Scientific) and reactions contained advanced SYBR green qPCR mastermix (Wisent), 50 nM of forward primer, 300 nM of reverse primer and 50 ng of DNA in a final volume of 10 μl. Reactions were performed in triplicates in 384-well plates using CFX384 Touch Real-Time PCR Detection System (Bio-Rad). We used a primer set designed to target the C. *rodentium* specific espB gene (espB-F: 5′-ATGCCGCAGATGAGACAGTTG-3′ and espB-R: 5′-CGTCAGCAGCCTTTTCAGCTA-3′, Invitrogen) which were previously determined to show sensitive quantification of *C. rodentium*(83). The cycling conditions were: incubation at 95°C for 10 minutes, followed by 40 cycles of 15 seconds of denaturation at 95°C, and 1 minute of annealing/extension at 60°C. For absolute quantification of bacterial loads, a standard curve was constructed using 10-fold serial dilutions of DNA extracted from a known number of *C. rodentium* cells, counted in triplicates using a hemacytometer.

### Detection of *C. rodentium* in tissue

To evaluate bacterial dissemination into distant tissues, liver and spleen samples were homogenized in 1 ml of sterile PBS, and serial dilutions were plated on LB agar plates with streptomycin and incubated aerobically at 37°C overnight. The number of colonies were counted and plotted as a CFU count/gram tissue.

### Histology

Colons were fixed in 10% formalin and were processed for paraffin-wax embedding using an automated processor. Tissue blocks were cut at 5 μm thickness, and sections were stained with hematoxylin and eosin (H&E). Digital light microscopic images were acquired using Aperio AT2 (Leica Biosystems, Ontario, Canada). A blinded experienced pathologist (A.G.) assessed the severity of colonic inflammation according to an established scoring scheme(84). Using the ImageJ Software(85), the length of at least 30 crypts were measured per mouse and a mean per mouse was calculated and analysed. Immunofluorescence and data analysis was performed for Ki67 as described previously(5). Imaging of Ki67 was performed on a Zeiss LSM980 with Airyscan 2 confocal microscope, using a 63x/1.4 NA Plan-APOCRAMAT oil immersion objective. Briefly, the average of four intestinal crypts for each sample was counted to calculate the mean for Ki67-positive cells per crypt.

### Serum Collection & ELISA for IL-22

Whole blood collected through cardiac puncture was allowed to clot at room temperature for 30 min. Samples were then centrifuged at 15,000 r.p.m., 4℃ for 20 min. Supernatants were transferred to 1.5 mL Eppendorf tubes and stored at −80 °C for downstream analysis. To quantify IL-22, serum was collected as described and an IL-22 ELISA (IL-22, Cat# ab223857) was performed according to the manufacturers’ instructions. IL-22 concentration was measured at 450 nm with a plate reader.

### Western Blotting

Colonic tissues were sonicated on ice in tissue lysis buffer (Thermo Scientific) with a cocktail of protease inhibitors (Sigma). Protein concentration was measured using BCA Protein Assay Kit (Thermo Scientific) as per manufacturer’s protocol. 20 μg of protein was electrophoresed through layered gels made in-house (polyacrylamide concentration: stacking gel, 4%; top resolving gel 12% and bottom resolving gel 16%). Trans-Blot Turbo Transfer System (Bio-Rad) was used for semi-dry transfer (14 min) onto a polyvinylidene fluoride (PVDF) membrane (Millipore) with 0.45 μm pore size. Membranes were blocked for 1 h at room-temperature in TBS-Tween 0.1% containing 3% BSA. Membranes were incubated in primary antibody overnight at 4 °C, followed by secondary HRP-conjugated antibody incubation for 2 hours at room temperature. Antibody concentrations used and catalogue numbers are detailed in Supplementary Table 3. Proteins were visualized using a Pierce enhanced chemiluminescence (ECL) kit (Invitrogen, USA), images were captured using the Odyssey Imaging System (LI-COR) and analyzed using Image Studio Lite v.5.2.5 (LI-COR). After quantification, Restore^TM^ Western Blot Stripping Buffer (Thermo Fisher Scientific, USA) was used to remove primary and secondary antibodies, allowing the same membrane to be reprobed with a different antibody. Uncropped raw blots are provided as Supplementary Figure 3.

### Statistical Analysis

Statistical analysis was done using Prism 9 software (GraphPad Software, San Diego, California USA). Data are represented as the mean ± SD. As detailed in figure legends, statistical comparisons and continuous normally distributed variables were evaluated with either unpaired, two-tailed student’s t-test (for two groups) or one-way ANOVA (for multiple groups) followed by Tukey’s multiple comparisons test. For ordinal, non-normal or categorical data, non-parametric unpaired, two-tailed Mann-Whitney test, and Fisher’s exact test was applied. Linear mixed-effect model was applied for the clinical severity score data. Kaplan-Meier survival was analyzed by Mantel-Cox Logrank test. To assess group differences in repeated measures, two-way ANOVA followed by Tukey’s multiple comparisons test was used. P <0.05 was considered statistically significant.

## Acknowledgements

The authors thank the Imaging Facility at The Hospital for Sick Children for their technical help and support. We also want to thank Bernald Castro for technical support with histological staining. This work would not have been possible without Joel Tan (Dr. John Brumell lab), and Sharon Leung and Emiliano Miraglia (Dr. Peter Kim lab), who provided the wildtype strains of the bacterial pathogens and helped with the microbiology setup in the lab. This research was funded by the Bill & Melinda Gates Foundation (OPP1185057) and the Canadian Institutes of Health Research (CIHR156307).

## Author contributions

M.K.T., M.F., and C.B. designed the study, performed the experiments, and analyzed the data. S.D.W. performed the experiments for Figure 1. J.P., C.C., and E.M.C. helped with the qPCR set-up for Figure 2a. A.G. scored histological results for Figure 3. S.L. produced the Ki67 immunofluorescent images for Figure 4. M.K.T., M.F., C.B., A.F., and R.H.J.B. interpreted the results. M.K.T wrote the first draft and prepared the figures. All authors contributed to editing and critical review. M.K.T., M.F., C.B., A.F. and R.H.J.B. conceived the project. A.F. and R.H.J.B. supervised and coordinated the work and R.H.J.B. secured project funding.

## Data availability

Data will be made available from the corresponding author on reasonable request.

## Additional Information

**Supplementary Information** The online version contains supplementary material available at xx

